# Epigenetic and evolutionary features of ape subterminal heterochromatin

**DOI:** 10.1101/2025.05.29.656835

**Authors:** DongAhn Yoo, Katherine M. Munson, Evan E. Eichler

**Affiliations:** Department of Genome Sciences, University of Washington School of Medicine, Seattle, WA, USA; Howard Hughes Medical Institute, University of Washington, Seattle, WA 98195, USA

**Keywords:** Genomics, Subterminal heterochromatin, Evolution, Great ape, Comparative genomics

## Abstract

Many African great ape chromosomes possess large subterminal heterochromatic caps at their telomeres that are conspicuously absent from the human lineage. Leveraging the complete sequences of great ape genomes, we characterize the organization of subterminal caps and reconstruct the evolutionary history of these regions in chimpanzees and gorillas. Detailed analyses of the pCht satellite composition and associated segmental duplication (SD) spacers confirm two independent origins in the *Pan* and gorilla lineages. In chimpanzee and bonobo, we estimate these structures emerged ∼7.5 million years ago (MYA) in contrast to gorilla where they expanded more recently ∼5.1 MYA and now make up 8.5% of the total gorilla genome. In both lineages, the SD spacers punctuating the pCht heterochromatic satellite arrays correspond to pockets of hypomethylation, although in gorilla such regions are significantly more hypomethylated (*p*<2.2e-16) than chimpanzee or bonobo. Allelic pairs of subterminal caps show a high degree of sequence divergence (9-11%) with bonobo showing less divergent haplotypes and less hypomethylated spacers. In contrast, we identify virtually identical subterminal caps mapping to nonhomologous chromosomes within a species, suggesting ectopic recombination potentially mediated by SD spacers. We find that the transition regions from heterochromatic subterminal caps to euchromatin are enriched for structural variant insertions and lineage-specific duplicated genes. We suggest these regions are hotspots for accelerating ape genome evolution.

## INTRODUCTION

With respect to humans, chromosome karyotypes of nonhuman African great apes (chimpanzee, bonobo and gorilla) differ by the presence of subterminal heterochromatic caps, which were recognized cytogenetically more than 40 years ago (Yunis and Prakash 1982). Among chimpanzee, gorilla and bonobo, the subterminal caps are differentially distributed among chromosomes. These caps are composed of hundreds of kilobase pairs (kbp) of long satellite arrays where the basic repeat unit, known as pCht satellite (also called as StSat or subterminal satellite), is 32 bp in length (Royle et al. 1994; Koga et al. 2011; Ventura et al. 2012). Previous cytogenetic studies in chimpanzees have shown that subterminal caps can form unique terminal associations in 32% of spermatocytes during late meiotic prophase I, so called post-bouquet structures. These cytogenetic structures are thought to promote ectopic recombination through persistent interactions between subtelomeric sequences among homologous and nonhomologous chromosomes (Hirai et al. 2019). The function of the subterminal caps is not known though have been proposed to help stabilize African great ape genomes by preventing or buffering against interchromosomal exchanges of more proximal subtelomeric sequences (Hirai et al. 2019) or contributing to the replication biology of telomeres (Novo et al. 2013).

Subsequent investigations into the organization of these regions discovered segmental duplication (SD) “spacers” interdigitated between the large arrays of pCht satellite DNA (Ventura et al. 2012; Yoo et al. 2025). SD spacers of different phylogenetic origin were identified in *Pan* and gorilla lineages suggesting that the subterminal cap structures arose independently. Ventura et al. proposed an evolutionary model, where a human–chimpanzee pericentric inversion followed by chromosome 2 fusion contributed to the predisposition or loss of subterminal caps in nonhuman apes and human genomes (Ventura et al. 2012). Jiang and colleagues further refined the chromosome 2 fusion breakpoint and subterminal SDs in humans (Jiang et al. 2024). In the most recent study, the heterochromatic cap sequences were fully resolved using a hybrid long-read assembly approach (ONT and HiFi) and assembly phasing (HiC and parent–child trios) (Yoo et al. 2025).

While these subterminal satellites are absent in other apes like orangutans (Haaf and Schmid 1987; Ijdo et al. 1991; Ventura et al. 2011), such structures are not unique to African great apes. A similar, albeit larger, subterminal heterochromatic cap structure composed of alpha-satellite DNA was described for siamang genomes from the gibbon lineage (Koga et al. 2012). The independent emergence of subterminal satellite structures in multiple primate lineages (gorilla, chimpanzee and siamang) suggests convergent evolution of potential functional significance. In this study, we leverage the fully resolved sequence of telomere-to-telomere ape genomes to more systematically investigate the evolution, structure, and epigenetic properties. The availability of two fully resolved haplotype assemblies from each ape species allowed for investigations into patterns of allelic variation and between-species comparisons of subterminal organization and differences in methylation patterns. We examined the telomeric transition regions from heterochromatin to euchromatin leading to potentially new insights into the genic content and stability of these regions across ape genomes.

## RESULTS

### African great ape subterminal heterochromatic cap structure and methylation

Using the recently published ape telomere-to-telomere (T2T) genomes (Yoo et al. 2025) (**Supplementary table 1**), we estimated the size and content of each chromosomal cap in chimpanzee, bonobo, and gorilla (**Fig. 1 & Supplementary table 2**) defining chromosomes based on their synteny with human (McConkey 2004). Overall, gorilla chromosomes have larger and greater number of subterminal caps where they account for 584.1 Mbp (8.46%) of the diploid genome, while that of chimpanzee and bonobo genomes represent 337.9 and 314.7 Mbp (5.4% and 5.0%), respectively. We identified subterminal caps on both homologs (allelic pairs) in both chimpanzee and gorilla, while bonobo showed patterns consistent with heteromorphic variation (present on only one of two homologous chromosomes). For example, the bonobo p-arms of chr14 (hsa13) and chr19 (hsa17), and both p- and q-arms of chr18 (hsa16) possessed subterminal caps on only one of the two haplotypes for this individual. This heteromorphism and the fact that bonobo possesses the lowest number of subterminal caps (n=46 compared to 57 in chimpanzees and 79 in gorillas) may suggest their subterminal caps could be in a state of decay—gradually being lost from the bonobo lineage. Notably, the short (p-) arms of chimpanzee and bonobo more consistently harbor a greater proportion of subterminal caps (39 and 31) when compared to q-arms (14 and 11). In contrast, the gorilla genome shows a more uniform distribution with more q-arm (45) subterminal caps than p-arms (34). The lengths of the subterminal caps were highly variable ranging from <1 Mbp in chimpanzee chr14 (hsa13), to a 35-fold longer length of 18.9 Mbp in bonobo chr6 (hsa7) (**Fig. 1a**). Unlike gorilla, two interstitial satellite arrays are observed in chimpanzee and bonobo mapping to chr6 (hsa7) and 14 (hsa13).

**Figure 1.**
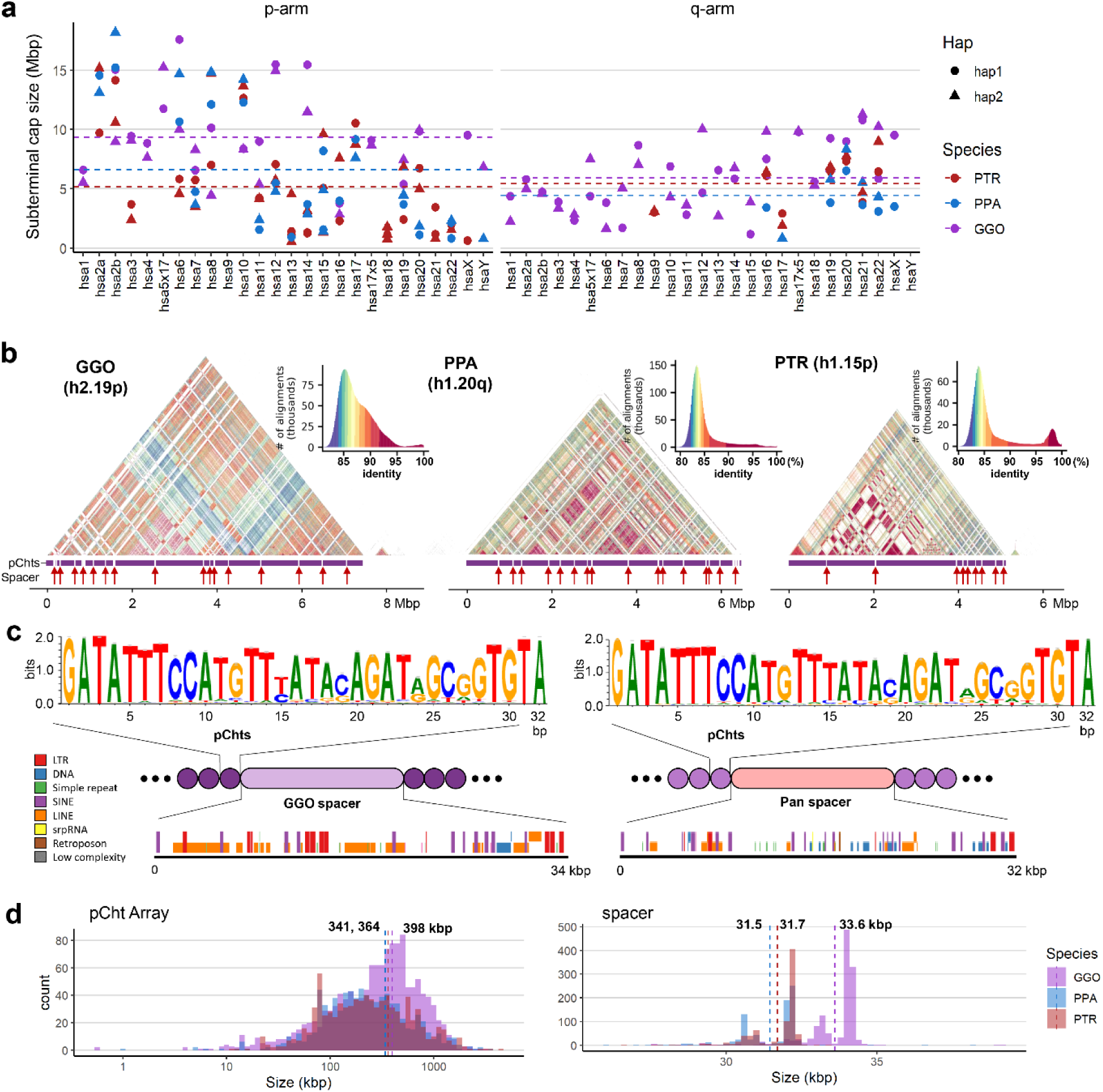
Overview of African great ape subterminal cap organization. a) Size of p- or q-arm subterminal caps in three species: chimpanzee (PTR), bonobo (PPA) and gorilla (GGO), indicated by red, blue and purple, respectively. Circles and triangles indicate haplotypes 1 and 2, or maternal and paternal for bonobo and gorilla, respectively. b) StainedGlass (Vollger et al. 2022) self-alignment plot of subterminal caps including pCht satellites indicated by the purple track below the triangular heatmap, as well as subterminal SD spacers interrupting satellite arrays, indicated by the red arrows below. c) The structure of subterminal caps. From the top, the logo base profile of pCht satellite is shown, followed by the higher-order structure of subterminal cap sequences, and the sequence composition of the subterminal SDs at the bottom. d) Size distribution of uninterrupted pCht array and subterminal SDs. The dotted line indicates the mean. The x-axis is in a log10 scale for the pCht array due to variation in its size.

As previously reported (Yoo et al. 2025), analysis of the subterminal caps revealed a higher-order organization of pCht satellite arrays of variable length (ranging from <10 kbp to hundreds of kbp with a mean of 341-398 kbp) interdigitated with SD spacers (**Fig. 1b-d**). We find that pCht satellites are generally more than 80% identical with the ones in closer proximity showing higher identity within a chromosomal arm (**Fig. 1b**). While different subtypes of pCht can be distinguished, we derived a consensus for each depending on species (**Fig. 1c & Supplementary fig. 1**). Despite the sequence variation, we find that the gorilla satellite consensus is almost identical to that of the *Pan* lineage (chimpanzee and bonobo), with the exception of the thymine in the 15^th^ position, which shows less variability in the *Pan* lineage. In terms of epigenetic status, these satellite sequences show high levels of 5mC methylation, with gorilla showing average methylation of 83.3% compared to 78.4% in bonobo and 73.6% in chimpanzee (**Supplementary figs. 2 & 3a**).

The pCht satellite arrays are interrupted irregularly by hypomethylated pockets of subterminal SD spacers (**Supplementary figs. 2 & 3**). These SDs originate from euchromatic sequence in the ancestral ape lineage (**Fig. 1c**). For example, the ancestral subterminal SD spacer in gorilla maps to the intronic region of the gene *MALD1* of chr8 (hsa10, p12.31) and subsequently expanded to ∼700 copies per gorilla haplotype. In contrast, the spacer of the *Pan* lineage originates from a subsequence of *PGM5*, chr11 (hsa9, q21.11) (**Supplementary fig. 4**), and expanded to ∼413 and ∼430 copies per haplotype for chimpanzee and bonobo, respectively (**Supplementary table 3**). The average length of gorilla and *Pan* spacers are 34 and 32 kbp, respectively (**Fig. 1c-d**). Of note, the length of both spacers and pCht arrays are significantly longer in gorilla compared to chimpanzee and bonobo (two-sided Wilcox test *p*<2.2e-16 and *p*=6e-10, respectively) consistent with their independent origin and expansion in each lineage (Ventura et al. 2012).

Examining the epigenetic status of SD spacers, we find a significantly lower rate of 5mC methylation in gorilla (average of 38.8%) relative to chimpanzee (40.2%) and bonobo (49.9%), with *p*<2.2e-16, two-sided Wilcox test (**Supplementary figs. 2 & 3b**). Using SD spacers mapping outside of pCht satellite arrays as a comparative control, we find that 99.8% and 98.5% of gorilla and chimpanzee spacers are more hypomethylated while only 17.8% of bonobo spacers are more hypomethylated than their interstitial paralogs (**Supplementary fig. 3b**). In chimpanzee and bonobo, spacer length positively correlates with CpG methylation although no such pattern is observed for gorilla where spacer hypomethylation is much more uniform (**Supplementary Fig. 3c**). We searched for evidence of transcription using Iso-Seq data previously generated for these individuals (Makova et al. 2024; Yoo et al. 2025) and found no credible evidence of novel spliced transcripts originating from the pCht satellite arrays or SD spacers as opposed to a previous study (Novo et al. 2013) (**Supplementary tables 4-5**).

### Evolution of the spacer and satellite sequence

In order to reconstruct the evolutionary history of the subterminal satellites, we performed an independent analysis of the two classes of repeats which define its composition: the pCht (∼32 bp) satellite that exists in millions of copies throughout the ape genomes and the *Pan* and gorilla SD spacers (32-34 kbp) that represent independent SD expansions (400-700 copies per genome) in association with the pCht satellites. While the SD spacer is far less abundant than the pChts, its length is thousands of times longer, providing a robust signal for multiple sequence alignment (MSA), phylogenetic reconstruction, and timing estimates.

For the pCht satellites, we first extracted all pCht units from chimpanzee (n=6,881,426), bonobo (n=6,616,396), and gorilla (n=14,248,529) subterminal caps. From this, we selected the most abundant 19,012 pCht variants (85% of the total pChts) that occur more than 100 times in these genomes. Under the assumption that more closely related subterminal arms would likely share a similar pCht composition, we performed an all-to-all pairwise comparison between subterminal heterochromatic caps that contain at least 2,000 pChts to identify those that are more closely related to one another. We applied unsupervised hierarchical clustering (pvclust) (Suzuki and Shimodaira 2006) across the three species and constructed a tree based on normalized counts of each pCht variant between subterminal heterochromatic caps (**Fig. 2a, Methods**). The composition of pCht variant types clearly distinguishes the gorilla and *Pan* lineages with the exception of chr5 (hsa6) q-arms of gorilla, which groups with the subterminal caps (chrX) from the *Pan* lineage (**Fig. 2a**). This suggests a possible common origin, but the majority of the subterminal caps have formed independently as a result of recurrent expansions in each lineage. Among the subterminal caps from chimpanzee and bonobo, pCht variant compositions do not distinguish between the two *Pan* species indicating that this expansion largely occurred prior to the speciation of bonobo and the common chimpanzee.

**Figure 2.**
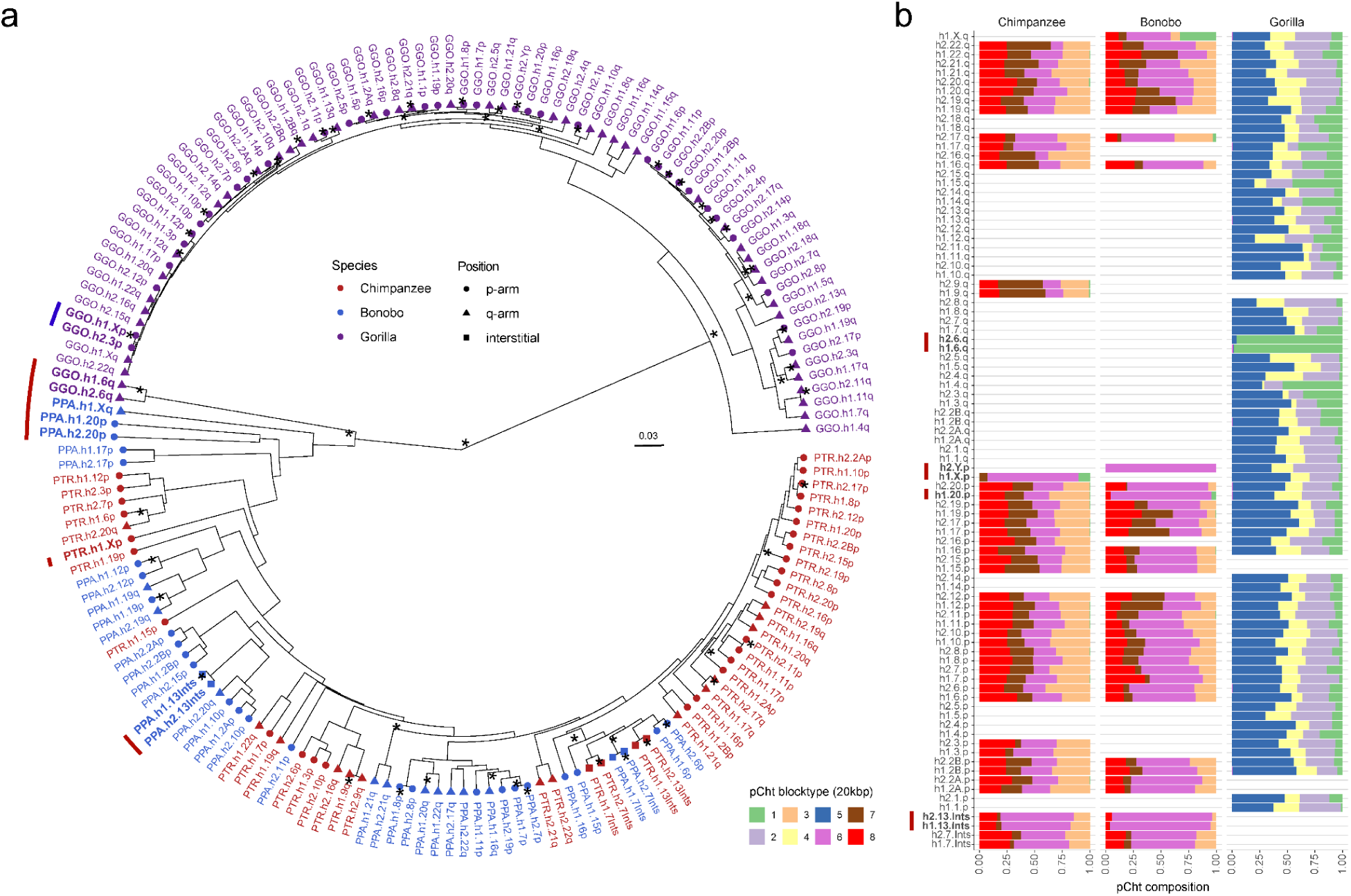
pCht satellite higher-order structure and phylogeny. a) Clustering of the subterminal arms based on relative abundance of pCht variants. Each terminal node indicates species (chimpanzee - PTR, bonobo – PPA, and gorilla - GGO), human chromosome number, and whether the position of the subterminal cap is located in p-arms, q-arms, or interstitial (Ints). The top 5% of subterminal caps with the largest proportion of shared pCht types (among GGO and *Pan*) are highlighted (bold/red line) (**Supplementary fig. 6c**). GGO.h2.3p vs. GGO.h1.Xp (blue line) is an example where the non-allelic subterminal cap pair is more closely related than the allelic pair. Internal nodes with bootstrap score higher than 95 are indicated by ‘*’. b) Identification of eight higher-order block types (20 kbp) of pCht satellites across the subterminal caps of three African great apes. The highest proportion of shared pCht types among chimpanzee and gorilla are indicated (red line).

**Figure 3.**
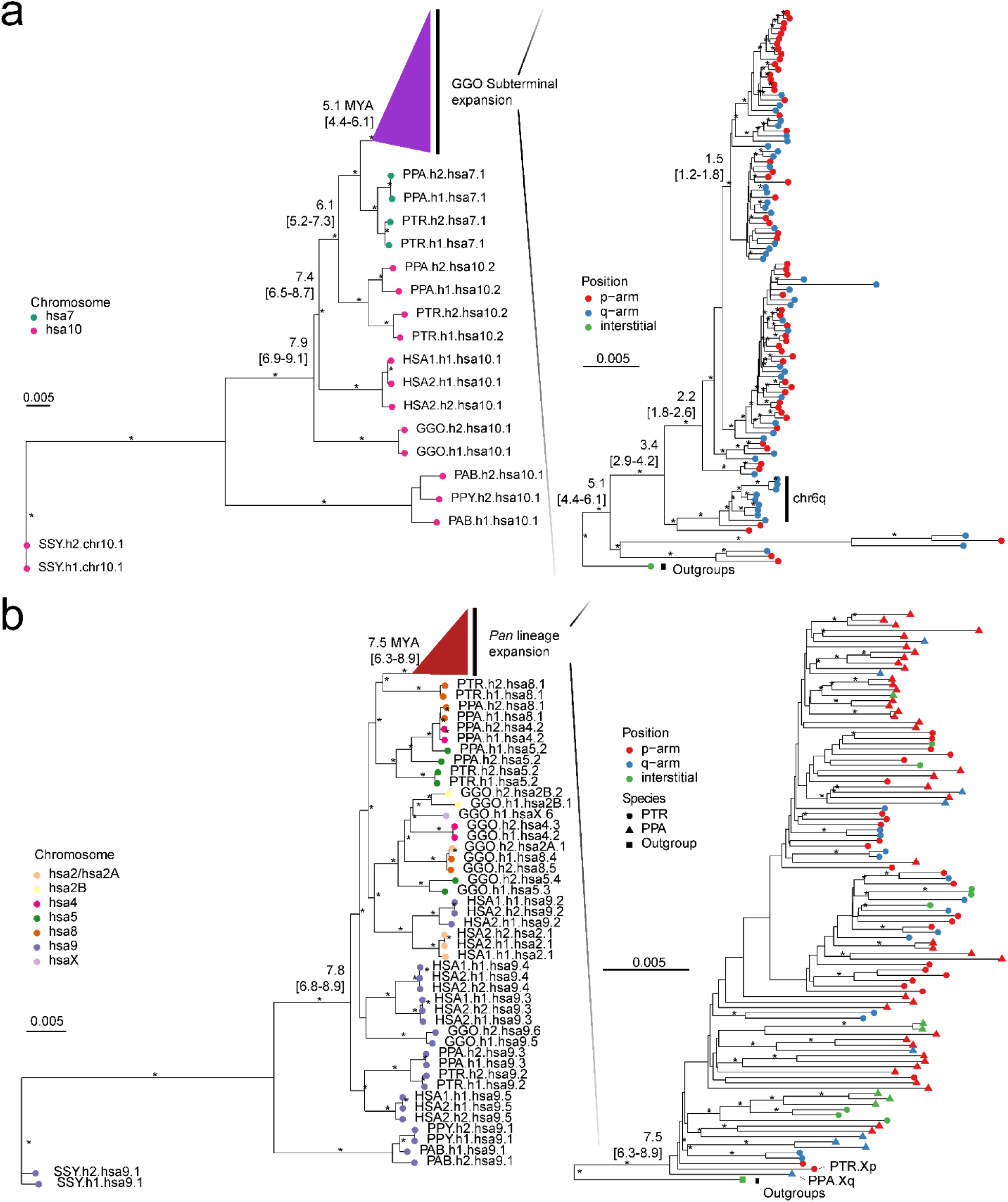
Evolution of subterminal SD spacers. a) Maximum likelihood phylogenetic tree of gorilla (34 kbp) and b) *Pan* subterminal SDs (32 kbp) constructed from a representative subset of spacers (n=100 copies). In each, the left panel shows the overall topology with a time-scale tree calibrated to orthologous ancestral primate copies. Chromosome numbers (based on human chromosome synteny) are color-coded and labelled by haplotype (h1 or h2) and species acronym: human (HSA; HSA1-CHM13 and HSA2-HG002), chimpanzee (PTR), bonobo (PPA), gorilla (GGO), Sumatran orangutan (PAB), Bornean orangutan (PPY), and siamang (SSY). A colored triangle denotes the subterminal spacer expansion. The right panel zooms into the subterminal spacer topology and time estimates in million years ago (MYA) with 95% confidence interval in the bracket. Gorilla (top) and chimpanzee (bottom) spacers are classified as p-arm (red), q-arm (blue), or interstitial (green). In addition, for the bottom panel, species are indicated as outgroup (rectangle), chimpanzee (circle), or bonobo (triangle) in origin. Internal nodes with bootstrap score higher than 95 are indicated by ‘*’.

Intraspecies sequence comparisons of the subterminal satellites show that some nonhomologous chromosomes are more similar in composition than their allelic counterparts—i.e., homologous subterminal caps do not always pair (e.g., GGO.h2.3p vs. GGO.h1.Xp; **Fig. 2a**). Under the assumption that nonhomologous subterminal caps are subject to ectopic recombination (Ventura et al. 2012; Hirai et al. 2019), we sought to classify the major pCht subtypes using a two-step k-means clustering approach (**Methods, Supplementary fig. 5**). We identified eight higher-order satellite block types across the 167 subterminal caps (**Fig. 2b**) from the three ape species.

Consistent with the hierarchical clustering tree, the gorilla and *Pan* lineages are largely distinct. Gorilla subterminal caps consist of k2, k4, and k5 block types while the *Pan* lineage caps are composed of k3, k6, k7, and k8 types. Once again, gorilla chromosome 5 (hsa6) is an exception with a largely uniform composition of the k1 block type, which is shared with the X chromosome p-arms of bonobo and chimpanzee and to a lesser degree with a fraction of other *Pan* autosomes. The k1 block type, thus, is distinct in being the only block type shared among the gorilla and *Pan* lineages, potentially identifying the ancestral pCht. We also note that this block type is composed of the largest portions of common pChts shared among *Pan* and gorilla (**Supplementary fig. 6**).

As a second approach, we used the SD spacer sequences of the gorilla and *Pan* lineages as a proxy to trace the evolutionary history of the subterminal heterochromatic caps. In this approach, we extracted a shared segment (>20 kbp in length and >90% identity) from the spacer for gorilla and the spacer for the *Pan* lineage and generated an MSA to construct a maximum likelihood phylogenetic tree for each lineage (**Methods & Fig. 3**). Using Asian apes as an outgroup (**Methods**), the gorilla spacer phylogenetic tree predicts that the ape ancestral locus (hsa10) began to duplicate 6.1 (5.2-7.3) million years ago (MYA) in the common ancestor of gorilla and chimpanzee (but not human) creating an interstitial copy mapping to *Pan* chr6 (hsa7) that is located 2 kbp away from an interstitial pCht satellite array. This interstitial spacer in the *Pan* lineage is ancestral to all other gorilla subterminal cap spacers beginning to duplicate soon thereafter (5.1 [4.4-6.1] MYA) (**Fig. 3a**), although it is no longer identified at the syntenic location in gorilla or human. The topology of the gorilla phylogenetic tree suggests a series of initial stepwise duplications, e.g., the gorilla chr5 (hsa6) q-arm is estimated to have occurred 3.4 (2.9-4.2) MYA. After these initial duplications, a subsequent burst of gorilla spacers began to occur 1.5 to 2.2 MYA giving rise to the majority of SD spacers in the gorilla genome. We also note that the phylogenetic tree of the duplicated spacers shows signs of incomplete lineage sorting, which does not follow the species tree (**Fig. 3**).

Although the SD spacer in the *Pan* lineage differs with respect to its ancestral origin (hsa9), similar to the gorilla it is predicted to begin its duplication in the common ancestor of humans, chimpanzees, and gorillas creating multiple interstitial copies (hsa4, 5, 8, and X) as well as copies located subterminal of hsa2A/2B—the fusion point (Jiang et al. 2024) for the formation of human chromosome 2 (**Fig. 3b**). Overall, branch lengths or sequence identity of the chimpanzee phylogeny are significantly longer (*p*<2.2e-16 by two-sided Wilcox test; **Supplementary fig. 7**) than gorilla consistent with a series of more ancient duplications in the *Pan* lineage. We estimate the oldest divergence time among the *Pan* lineage SD spacers to be 7.5 (6.3-8.9) MYA, which is close to the divergence time of human and gorilla ancestral orthologs (7.8 MYA). The chimpanzee SD spacers associated with the chrX p-arm subterminal caps, which also share ancient higher-order pCht blocks, represent some of the most ancient spacer duplications (**Fig. 3b**). Similar to gorilla, the topology of the chimpanzee spacer tree predicts a burst of duplications, albeit much older (3.6-3.8 MYA—average of terminal branch length) when compared to gorilla (0.6 MYA; **Supplementary fig. 7b**). It should be noted, however, that the spacers from homologous chromosome arms do not always cluster, potentially as a result of ectopic recombination (**Supplementary fig. 8**).

### Patterns of allelic and non-allelic variation

A striking feature of the subterminal heterochromatic caps is the high degree of allelic sequence diversity observed between two haplotypes (**Fig. 4a-c; Supplementary data**). Only 12% (10/82) of haplotype comparisons show >99% sequence identity with >50% alignment coverage. Most haplotype comparisons show greater than 10% sequence divergence (**Supplementary figs. 9-12**). A comparison with other genomic regions, including centromere, acrocentric and remaining regions, reveals that subterminal caps were the regions with the greatest degree of allelic divergence (**Supplementary fig. 13**). Notably, the degree of allelic divergence differs among the species. Gorilla and chimpanzee show on average greater sequence divergence (10.8 and 11%, respectively) when compared to bonobo (8.7%) (**Fig. 4d**). When comparing non-allelic versus allelic sequence identity, gorilla and chimpanzee distributions are largely indistinguishable, in contrast to bonobo where allelic alignments now show significantly greater allelic sequence identity when compared to non-allelic alignments (**Fig. 4e**). This feature is reflected in a matrix percent identity plot where each subterminal haplotype is aligned to every other within the individual (**Fig. 4b,c**). In contrast to bonobo where more than a third of allelic comparisons show the highest degree of sequence identity (as indicated by the diagonal in **Fig. 4c**), gorilla shows no such pattern.

**Figure 4.**
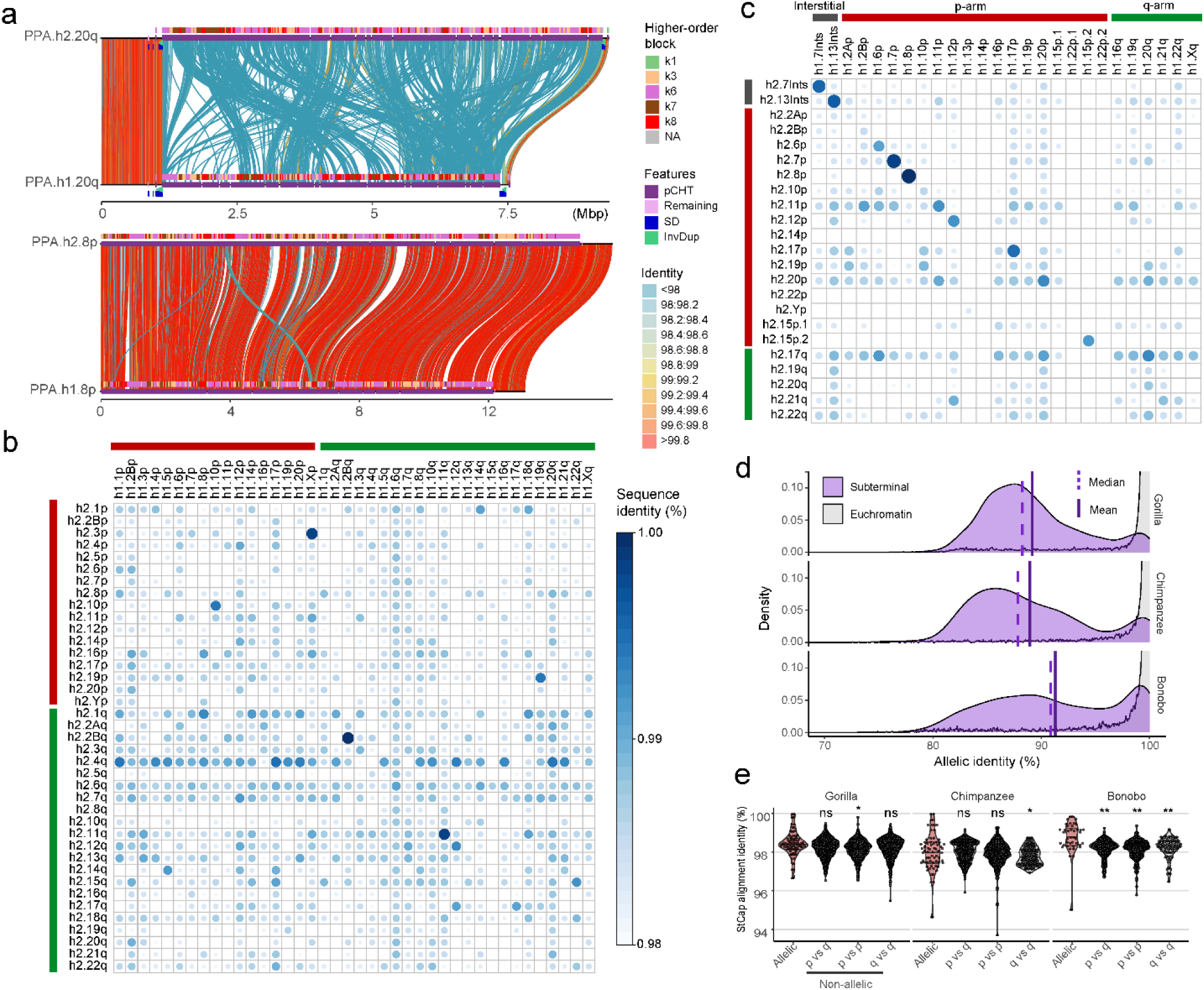
**Subterminal allelic vs. non-allelic sequence identity**. a) Example of low-identity (hsa20q) and high-identity (hsa8p) allelic subterminal caps in bonobo. Annotations include: percent identity of alignment (scaled from blue <98% to red >99.8%); higher-order block type of pCht satellite arrays (top), satellites (middle) and SD content (bottom). b,c) Sequence identity matrix between paternal (h2) and maternal (h1) haplotypes of b) gorilla and c) bonobo comparing allelic versus non-allelic subterminal loci. Circle (size) and intensity (blue) indicates higher sequence identity. d) Distribution of allelic sequence identity of 50 kbp non-overlapping windows of three African great ape genomes. Allelic sequence identity of subterminal caps (purple) and the euchromatic regions (gray; excluding centromere, acrocentric and subterminal regions) are compared with mean (straight) and median (dashed line) indicated. e) Distribution of average sequence identity between allelic subterminal caps compared to non- allelic subterminal caps. Each dot represents the pairwise comparison. Two-sided permutation test significance is indicated on top; **: *p*<0.0001 and *: *p*<0.05.

Instead, allelic and non-allelic patterns are largely indistinguishable perhaps providing evidence of more rapid, ongoing ectopic recombination occurring in this lineage.

As part of our all-by-all alignment analysis of subterminal caps, we searched for specific examples where there was evidence of a recent ectopic exchange (>99.5% identity) between nonhomologous chromosomes. We identified 11 candidates for ectopic exchange (8 in gorilla, 3 in chimpanzee, and none in bonobo). For example, we identified a potential 5 Mbp ectopic exchange between GGO.3p (chr2/hsa3) and chromosome Xp (**Fig. 5a**). We refined the breakpoints of the sequence exchange and mapped both to different SD-spacer regions— corresponding to the 3^rd^ and 25^th^ SD spacer positions on the short arm of the gorilla X chromosome (**Fig. 5b**). Analyzing the remaining candidates (**Fig. 5c; Supplementary figs. 14- 16**), we found 3/8 examples where the breakpoint mapped precisely within the spacer and two additional examples where one of the breakpoints mapped within 10 kbp of a spacer. Given that the SD spacers represent a small fraction of the subterminal heterochromatic cap, we performed a simulation to test the proximity of breakpoints to SD spaces. We find that candidate breakpoints of recent ectopic exchange (>99.8 identity) are more likely (*p*=0.035, one-sided Wilcox ranked sum test; **Supplementary fig. 17**) to map in close proximity to SD spacers than expected based on a null genomic distribution (**Fig. 5d**).

**Figure 5.**
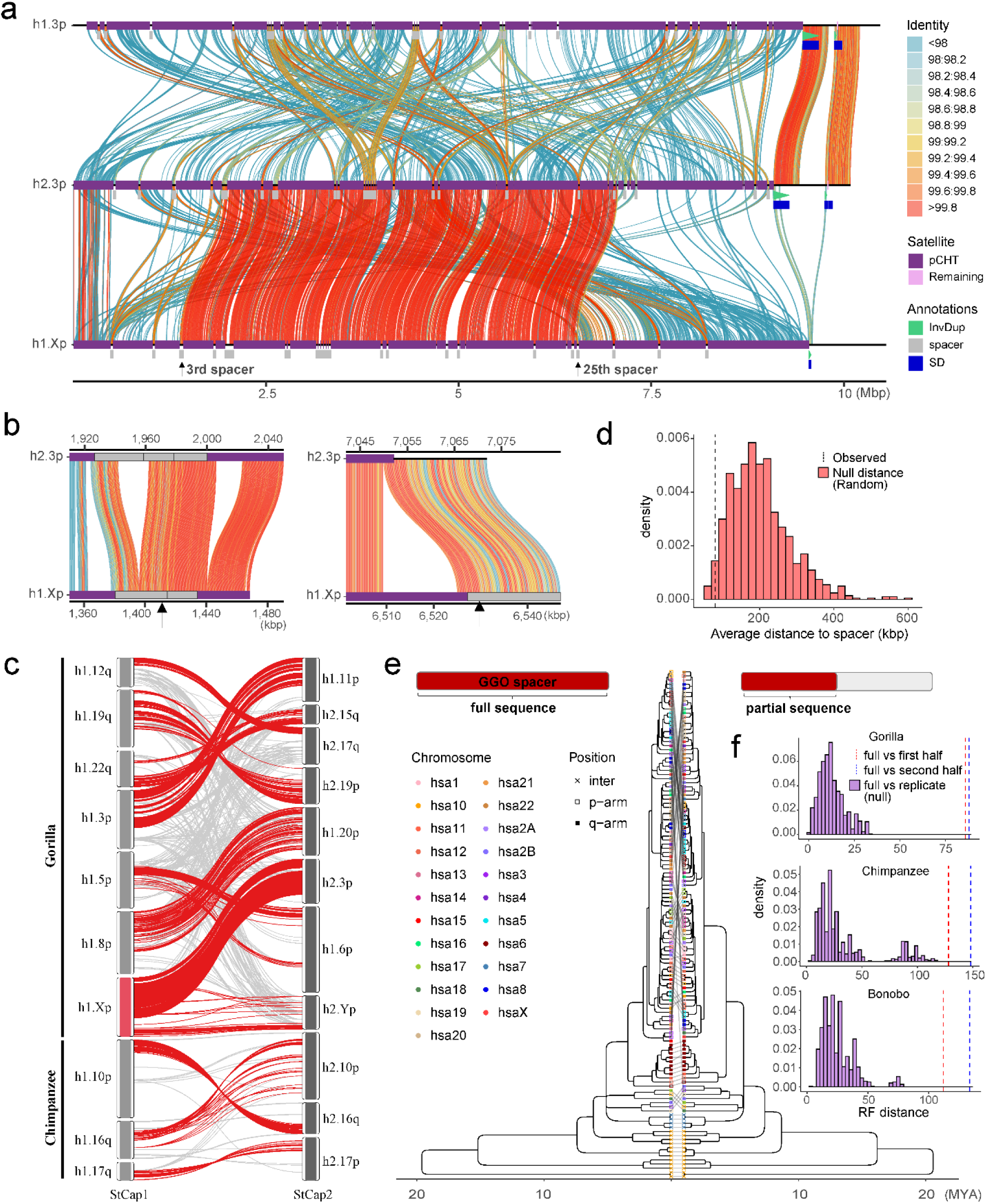
Evidence of non-allelic sequence exchange at SD spacers. a) An alignment view (SVbyEye)(Porubsky et al. 2024) depicting a candidate ectopic exchange between subterminal satellite chromosomes hsa3 and hsaX p-arms of gorilla. The percent identity of alignment (scaled from blue <98% to red >99.8%) with annotation tracks for higher-order pCht blocks, satellites, and SDs are shown on the right. b) Enlarged view of the two exchange breakpoints (black arrows) mapping within the 3^rd^ and 25^th^ SD spacers. c) Overview of potential ectopic exchange events between the subterminal caps (left-StCap1 vs right-StCap2 tracks) identified in African great ape genomes (eight and three events in gorilla and chimpanzee, respectively, including hsa3p vs. hsaXp in red). Assembly quality at the breakpoints was validated by a read-depth analysis in **Supplementary figs. 19-25**. d) Simulation test of exchange event break points, suggesting that the exchange breaks (>99.8% identity alignment blocks; dotted line) are more likely to be located within or close to subterminal SD spacers than a random distribution (red histogram). e and f) Comparison of maximum likelihood phylogeny to test recombination of SD spacer. A phylogenetic tree of the complete spacer sequence (left) is compared to that of a subset of the sequence (right). The tip nodes of the left tree are linked to the corresponding nodes of the right tree to visualize phylogenetic shifts in the topology. Robinson-Foulds distance is used to measure the extent of phylogenetic shift (first or last half sequences, in red and blue dotted lines and is compared to a null distribution generated by computing distances of consensus tree vs. bootstrap replicate trees).

As a final test of the potential role of SD spacers in mediating ectopic exchange, we developed a phylogenetic approach to test for this association. We reasoned that if spacers were preferential sites of ectopic recombination during evolution, dividing the ∼30 kbp spacer sequence into two segments (first and second half, located at telomere and centromere directions, respectively) would capture two distinct phylogenetic histories when comparing to each other (**Supplementary fig. 18**) or to the topology of the full SD spacer (**Fig. 5e**). To test for significance, we calculated the distribution of Robinson-Foulds metric between individual bootstrap tree topologies (each replicate with n=1000) to their consensus topology to generate a null distribution of the expected statistic and contrasted this with the partial sequence trees (**Fig. 5f**). We find a significant shift in the topology of the SD spacer subset trees (*p*<0.001) for all three species (gorilla, chimpanzee and bonobo), suggesting that historically the SD spacers may have been the preferential point of ectopic exchange between nonhomologous chromosome arms.

### Subterminal heterochromatic-euchromatic transition regions and the evolution of new genes

Finally, we examined the euchromatin boundaries and putative evolutionary consequences and epigenetic features of the subterminal caps (**Fig. 6**). Specifically, we compared ape species with (gorilla/chimpanzee) and without (Bornean and Sumatran orangutans) subterminal caps to determine if the rate of structural variation, gene density, and methylation differ in association with the evolution of these large terminal heterochromatic structures (Yoo et al. 2025). Using the human genome as a reference, we find ∼24–72 Mbp of novel insertions mapping within 2 Mbp from subterminal caps (African nonhuman apes) or telomeres (Asian great apes without subterminal caps) (**Fig. 6a-b**). The insertions in the African apes are notably more abundant (31.8–72.2 Mbp vs. 23.8–23.9 Mbp in orangutans) and larger (>10–500 kbp in size) when compared to the Asian great apes (**Fig. 6c** & **Supplementary fig. 26**). This is despite the fact that divergence time between humans to African great apes are close to half the divergence to Asian great apes (number of insertion base pairs scaled by time of divergence in **Supplementary fig. 27**). We estimate that 36%–42% of the inserted sequences in African apes originated from nonhomologous chromosomes (**Supplementary fig. 28**)—a potential consequence of the ectopic exchanges extending beyond the subterminal heterochromatic cap.

**Figure 6.**
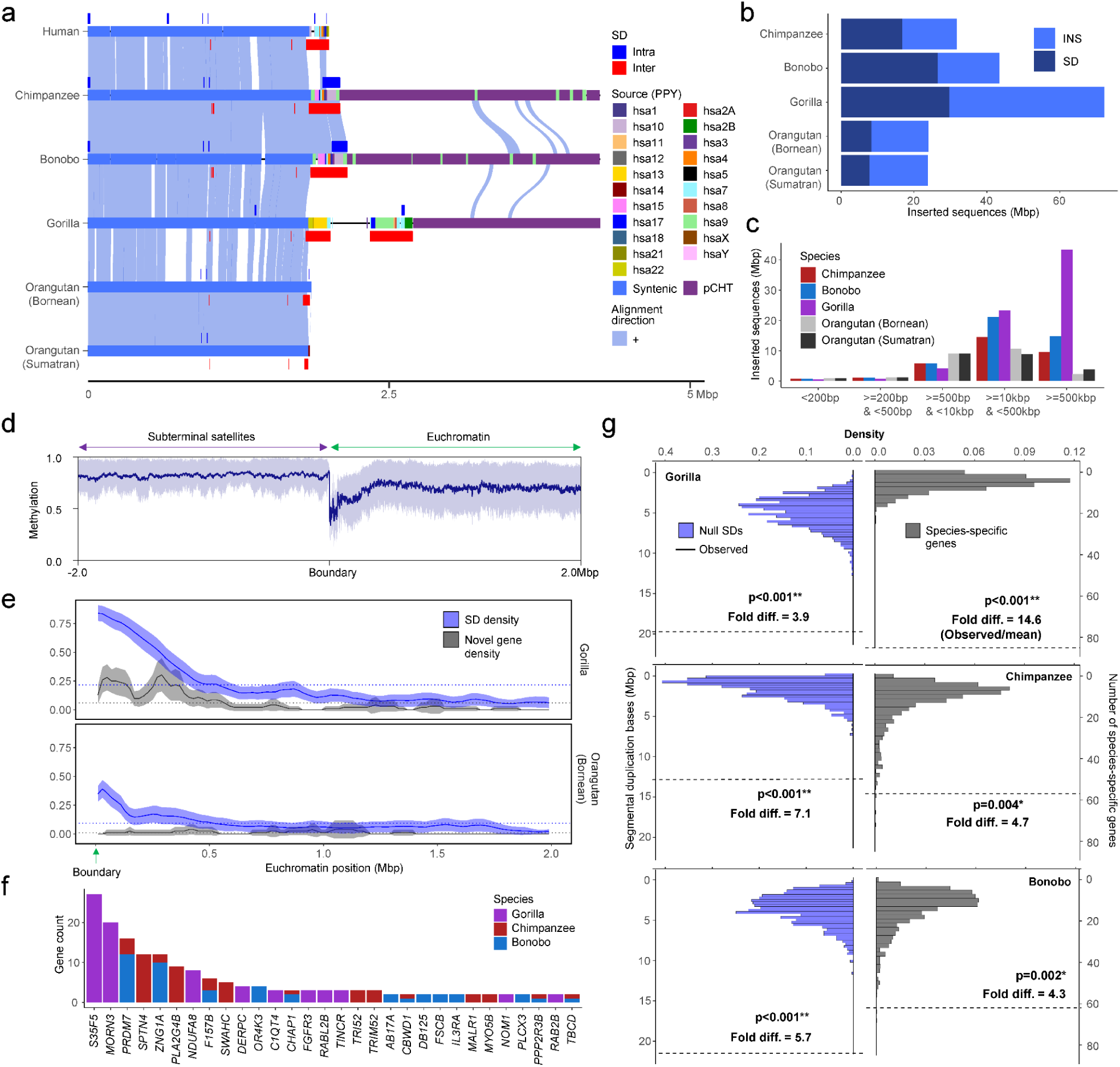
Duplicated genes and epigenetic features of ape genomes with and without subterminal caps. a) An ape comparative analysis of the organization of subterminal sequences for the long arm of chromosome hsa16. The stacked alignment plot (SVbyEye) contrasts syntenic euchromatic regions (light blue) with heterochromatic subterminal satellite regions (purple) and with the location of intra (dark blue) and interchromosomal (red) SDs along with the origin of the duplicated sequences (human chromosome designation). b) The total number of new sequences (insertions [INS] or SDs), not present in humans, at subterminal boundaries within 2 Mbp of the euchromatic-heterochromatic transition zone. c) Size distribution of the total inserted sequences with respect to human genome, color-coded by species. d) CpG methylation profile within 2 Mbp of the euchromatin-heterochromatin transition for chromosomes with subterminal caps in gorilla. The average % CpG methylation (blue line) with standard deviation (blue shaded area) is shown. e) The densities of SDs and novel genes at the euchromatic boundaries, highly enriched in gorilla, with subterminal caps, as opposed to orangutan. The transparent area indicates 95% interval of the observed density. Dotted horizontal line indicates the mean density. f) Duplicated genes and their copy number located at the boundary of subterminal caps, including eight genes validated by read depth of 120 primates (Mao et al. 2024) (**Supplementary fig. 35**). g) Simulations to test enrichment of novel genes (Yoo et al. 2025) and SDs at the boundary of subterminal caps (2 Mbp) in gray and blue, respectively. The *p* indicates empirical *p*-value of the one-sided permutation test.

Leveraging CpG methylation signals associated with the long-read sequence data used to generate these ape assemblies (Yoo et al. 2025), we investigated methylation at the euchromatin-heterochromatin boundary defined here as the last proximal subterminal pCht satellite unit.

Within 200 kbp of this transition, we observe an immediate and sharp drop (∼54%) in methylation among nonhuman African great apes (**Fig. 6d & Supplementary fig. 29 &** Asian great apes in **Supplementary fig. 30**), which quickly resets to a moderate level of hypermethylation (71%) when extended further euchromatically. Correlating with the transition to the hypomethylation, we also observe the highest density of SDs and species-specific annotated duplicated genes (Yoo et al. 2025), which also reduces when extended more proximally into the euchromatin of gorilla (**Fig. 6e**) and chimpanzee (**Supplementary fig. 31**). In Asian great apes, the signature of SDs and duplicated genes is less variable (**Fig. 6e** & **Supplementary fig. 31).**

In total we identify 204 novel duplicated genes at these subterminal euchromatin boundaries in gorilla (n=85), chimpanzee (n=57), and bonobo (n=62) (**Fig. 6f**). The enrichment of lineage-specific duplicated genes at this heterochromatin-euchromatin boundary is most extreme for gorilla (14.6-fold difference) followed by relatively high levels in the *Pan* lineage (4.3- to 4.7- fold) dropping to lower levels in the orangutan (1.6- to 2.8-fold) (**Fig. 6g & Supplementary fig. 32**). Compared to genome-wide average of each species, we find that the species with subterminal caps (nonhuman African great apes) have a greater enrichment of SDs (3.9- to 7.1- fold difference) than humans (3.0) or Asian great apes (2.3-2.5) at these boundaries (**Fig. 6g & Supplementary figs. 31 & 33**). In addition, we also find that the pairwise identity between the paralogous SD pairs at the euchromatin boundaries are the highest in gorilla when compared to other species (**Supplementary fig. 34**). In total, the result suggests an accelerated rate of evolutionary duplication and gene innovation among ape lineages possessing subterminal caps, potentially a consequence of both ectopic exchange and hypomethylation at the boundaries.

## DISCUSSION

The complete sequencing and analysis of the subterminal caps in both haplotypes of a single individual of chimpanzee, bonobo, and gorilla have provided novel insights into both the evolution and functional properties of these complex genomic regions. First, we provide supporting evidence for the largely independent expansion of the subterminal caps in both chimpanzee and gorilla as originally proposed by Ventura et al. (Ventura et al. 2012). Our phylogenetic analyses of both the satellite and the SD spacer sequences generally support this model with some important differences. The detailed pCht satellite higher-order structure analysis, for example, clearly defines a block type (k1) that likely existed in the common ancestor of both gorilla and chimpanzee lineages and we hypothesize that k1 was lost to the human lineage as a result of incomplete lineage sorting (ILS) or the chromosome 2 fusion (Jiang et al. 2024). Subsequent hyperexpansion of pCht in the gorilla and *Pan* lineages created lineage-specific higher-order block types that now account for the bulk of the subterminal heterochromatic caps of each species. Third, using the SD spacer sequences as a phylogenetic marker also supports early duplication and ILS. However, the phylogenetic analysis distinguishes two very different evolutionary trajectories. In the *Pan* lineage, both the formation (∼7.5 [6.3-8.9] MYA) and expansion (7.5-5.1 MYA) of the subterminal heterochromatin is much more ancient, far predating that of the gorilla SDs and the speciation of bonobo and the common chimpanzee. In contrast, gorilla subterminal heterochromatin began to emerge 5.1 (4.4-6.1) MYA—and hyperexpanded millions of year later (1.5-2.2 MYA). We predict the shared higher-order blocks (k1) of subterminal satellites identified in this study (**Fig. 2b**), as well as multi-copy orthologs of *Pan* lineage SDs in gorilla and human are the remnants of the ancient African great ape subterminal heterochromatin in these genomes.

While the general organization of the subterminal heterochromatic caps is similar among the nonhuman African apes, we document important epigenetic and genetic differences within each species. Owing to its more recent origin, gorilla subterminal heterochromatic caps are more homogenous among the nonhomologous chromosomes, and gorilla SD spacers show the most extensive CpG hypomethylation. In contrast, bonobo subterminal heterochromatic caps are composed of the oldest or most divergent subterminal SDs and are subject to heteromorphism (present on only one of the two allelic homologues) unlike other ape species (Ventura et al. 2012). Combined with a demonstrable weaker hypomethylation signal for the SD spacers (**Supplementary fig. 3**), much higher allelic identity (**Fig. 4**), and the fact that we find no cases of recent ectopic sequence exchange in bonobo genomes, this suggests that subterminal caps are in the process of decay although we caution that additional bonobo genomes derived from primary tissues as opposed to cell lines will need to be assayed to establish this as a species-specific property. Nevertheless, under this model, gorilla would be predicted to be the most active subterminal heterochromatic cap perhaps explaining why it is much more prevalent in this genome adding almost three chromosomes worth of DNA (584 Mbp).

Since the first characterization of African great ape subterminal satellites, there has been limited investigation into the functional significance of subterminal heterochromatic caps. Based on FISH experiments in chimpanzee meiotic sperm cells, a previous study by Hirai et al. 2019(Hirai et al. 2019) suggested that the subterminal heterochromatin was driving ectopic recombination by helping to form bouquet structures among nonhomologous chromosomes during late meiotic prophase somewhat akin to the nuclear organizing structures associated with the repeat-rich short arms of mammalian acrocentric chromosome (Nurk et al. 2022). Hirai and colleagues further concluded that such subterminal associations may be selectively beneficial by providing greater stability or buffering capacity to subterminal regions of ape chromosomes—areas known to be genetically and evolutionarily unstable (Knight and Flint 2000; Mefford and Trask 2002). In our study, we provide further support for their involvement in ectopic recombination by identifying 11 potential examples of interchromosomal translocations (8 in gorilla and 3 in the common chimpanzee; **Fig. 5c**). Both our breakpoint analysis of these events and our phylogenetic analysis indicate that the SD spacer regions are preferential drivers of this ectopic recombination between nonhomologous chromosome arms. It is possible that both the high degree of sequence identity of these 30 kbp segments and their hypomethylation status provide an ideal open-chromatin substrate for non-allelic homologous recombination. However, unlike Hirai’s prediction of conferring genomic stability, our evolutionary analysis suggests the complete opposite effect.

We find far stronger enrichment of insertions, especially interchromosomal SDs, near the euchromatin boundaries of chimpanzee, bonobo, and gorilla genomes when compared to Asian great apes without subterminal satellites (**Fig. 6**). More importantly, this enrichment has led to the emergence of 204 species-specific genes (Yoo et al. 2025) recently identified in chimpanzee, bonobo, and gorilla. These contain distinct splice junctions and transcript models with open reading frames and thus are candidates for the emergence of novel functional proteins in each species. Therefore, subterminal heterochromatic chromatin [similar to the pericentromeric and segmental duplications at the boundaries of inversions (Linardopoulou et al. 2005; Porubsky et al. 2020; Yoo et al. 2025)] may function as an incubator for the birth of new genes in ape genomes. This implicit genomic instability may confer a long-term selective advantage if the genes evolve function important for the survival of the species.

## METHODS

### Identification of subterminal satellites and SDs

Subterminal satellites or pCht repeats identified by a previous study were used (Yoo et al. 2025). Briefly, African great ape genomes (chimpanzee, bonobo and gorilla) were screened for pCht satellites via BLASTN (v2.12.0) (Chen et al. 2015) with the consensus sequence (len = 32 bp): “gatatttccatgtttatacagatagcggtgta”. BLAST hits with longer than 90% of the consensus (>28 bp) were recovered. The individual pCht unit was classified into different types based on the variants (small INS, DEL or substitution). The SD spacers interrupting the satellite arrays were identified using “bedtools subtract” option, subtracting the subterminal satellite arrays from the entire subterminal satellite regions, and 32 kbp of highly conserved sequence was identified in the *Pan* lineage and 34 kbp independently in gorilla. For extracting more precise locations of the individual SD spacers, minimap2 (Li 2018) (v2.26) was used (“*-x asm20*”).

The relationship between subterminal arms shown in Fig. 2a was predicted by performing hierarchical clustering implemented by pvclust v2.2 (Suzuki and Shimodaira 2006) analysis limiting to the most abundant pCht (n>100) (method.dist = “correlation”, and method.hclust = “average”, nboot = 100). The input matrix fed into pvclust was built by computing normalized counts of 19,012 pChts (count of one pCht variant in a subterminal arm/count of total pCht variants in that subterminal arm) across each subterminal arm. The higher-order structure of subterminal satellites was further investigated based on two-step k-means clustering (Hartigan and Wong 1979). The initial k-means clustering was performed under the assumption that some combinations of pCht variants are more frequently found together than others. This was done by computing the fraction of each pCht variant (n=19,012) across each of the subterminal arms (chr1p, chr1q, etc.). The optimal number of clusters was inferred by computing the Silhouette score (Rousseeuw 1987) for each k. The second k-means clustering was performed using the cluster of pChts (n=6) across subterminal caps divided into 20 kbp non-overlapping blocks.

Again, the optimal number of scores is determined with the Silhouette score. The second clustering identified a total of eight higher-order blocks of subterminal satellites that contain similar composition of pChts.

### Phylogeny of SD spacers

Based on the mutations accumulated in 32 and 34 kbp SD spacers, phylogeny was inferred using the maximum likelihood approach implemented by IQtree (Minh et al. 2020) (v2.3.6). A subset tree of 200 SD spacers were randomly sampled, and on top of the random set, the SD spacers located in chr6 q-arms were added to approximate their relative age. To estimate the position of the root, we identified ortholog copies of 32 and 34 kbp spacers in the remaining great ape lineages, using siamang as the outgroup; the ortholog copies were obtained with minimap2 (Li 2018) (v2.26). The MSA of the SD spacers was generated using MAFFT (Katoh and Standley 2013) (v7). Based on the MSA, two phylogenetic trees of SD spacers were constructed with -B 1,000, and with GTR+F+R6 and TVM+F+R5 models which were chosen as the best fit model via Bayesian information criterion, for gorilla and *Pan* spacer trees, respectively. The trees were time-scaled with the previously reported divergence to orangutan and siamang of 15.2 and 19.5 MYA, respectively (Kumar et al. 2022) (including IQtree parameters, “*--date-tip 0 --date-ci 100*”).

### Analysis of ectopic recombination

Under the assumption that ectopic sequence exchange takes place at subterminal arms, we investigated for evidence of such events via two different approaches, through pairwise comparisons of subterminal arms 1) restricted to SD spacers and 2) the full sequence. The alignment was performed independently in each species using minimap2 (Li 2018) (v2.26), and the average identity as well as breadth of coverage were computed. To locate the representative event, we narrowed down the search space, restricting to the alignment of identity >99.5% to obtain relatively recent events, and for the scale of sequences larger than 1 Mbp.

To investigate locational bias of ectopic sequence exchange breakpoints, we tested their relative distance to the nearest SD spacers. This was done through a simulation test, shuffling the breakpoints 1,000 times to generate the null distribution of average distances; the *p*-value was computed by quantifying the fraction of more extreme null values to the observed averages.

We further tested whether the breakpoints of ectopic sequence exchanges fall within SD spacers by computing phylogenetic trees. We divided the MSA of SD spacers into two equally sized regions and constructed the phylogenetic trees of the first and second half of sequences, which were then compared to the original tree computed by the full SD sequences. If the ectopic exchanges take place frequently at the SD spacers, the inferred phylogeny based on the first or second half of the sequence would shift dramatically. To test significance, we generated null distribution by computing the Robinson-Foulds distance between bootstrap trees to that of the consensus trees. This was then compared to the tree built by half of the SD sequences.

### Quantification of structural variations at the euchromatic boundaries

The subterminal euchromatic boundaries were defined as the 2 Mbp regions downstream or upstream of subterminal satellite arrays of p-arms or q-arms, respectively. In the case of species with no subterminal satellites, the boundaries were defined as the 2 Mbp tips. The structural variation calls from PAV (Ebert et al. 2021) (v2.3.2)—taking humans (T2T-CHM13 v2.0) as the reference and SDs identified from the previous study (Yoo et al. 2025)—were used. For the large insertions, we separately inferred the additional sequences of apes relative to humans by using syntenic alignment. The alignment between homologous chromosomes between human to nonhuman apes (i.e., chr1 vs. hsa1) generated by minimap2 (v2.26) were broken using RustyBam (v0.1.29) with a minimum structural event size of 50 bp. The unaligned regions of apes that are missing corresponding human homologs were considered as insertions. We also

performed alignment of individual chromosomes to the whole orangutan genome to investigate origins of the inserted sequences.

## DATA AVAILABILITY

The T2T genome assemblies generated by Yoo et al. 2025 (Yoo et al. 2025; **Supplementary table 1**) were used. In addition, the methylation track information available at https://github.com/marbl/T2T-Browser/tree/main/src/epi was used.

## ACKNOWLEDGMENTS

We would like to thank Tonia Brown, Soojin V. Yi, and Dongmin R. Son for helpful comments in the preparation of this manuscript. This work was supported, in part, by funding from the National Institutes of Health (NIH grants R01HG002385 and R01HG010169) to E.E.E. E.E.E. is an investigator of the Howard Hughes Medical Institute.

This article is subject to HHMI’s Open Access to Publications policy. HHMI lab heads have previously granted a nonexclusive CC BY 4.0 license to the public and a sublicensable license to HHMI in their research articles. Pursuant to those licenses, the author-accepted manuscript of this article can be made freely available under a CC BY 4.0 license immediately upon publication.

## AUTHOR CONTRIBUTIONS

D.Y. performed evolutionary and methylation analyses. D.Y. and E.E.E. wrote the manuscript.

K.M.M. provided sequencing support. E.E.E. and K.M.M. reviewed the manuscript. E.E.E. supervised and funded the project.

